# Conservation value of secondary and plantation forests for fruit-feeding butterflies in northern Vietnam

**DOI:** 10.64898/2026.10.07.757500

**Authors:** Thi Minh Thi Nguyen, Nhat Tan Pham, Van Lien Vu, Bao Thanh Le, Freerk Molleman

## Abstract

1. Assessing the conservation value of secondary forests and reforestation is important for conservation planning. Butterflies are widely used as indicators, and fruit-feeding butterflies are often used in biodiversity assessments in tropical landscapes.
2. We examined fruit-feeding butterfly diversity, abundance, and community composition across a primary forest, a secondary forest, and a plantation forest in northern Vietnam. We sampled butterflies nearly monthly during one year using baited traps.
3. Primary and secondary forest supported higher species richness than plantation forest. Moreover, while secondary forest had mainly a subset of species found in primary forest, the plantation forest had a distinct butterfly fauna.
4. Butterfly abundance and community composition varied across months, without showing a clear seasonal pattern.
5. Our results suggest that secondary and plantation forests do not fully replace primary forests, but they contribute meaningfully to butterfly conservation in human-modified tropical landscapes.

## INTRODUCTION

The loss and degradation of tropical forests are major drivers of biodiversity decline, making it increasingly important to evaluate the conservation value of secondary and planted forests (Barlow et al. 2007; Chazdon 2008; Gibson et al. 2011). While primary forests are irreplaceable for many forest-dependent species (Gibson et al. 2011), secondary forests may recover ecological functions over time through natural regeneration, and can then provide habitat, resources, and connectivity for many species (Chazdon 2008; Gibson et al. 2011; Korkiatupa et al. 2023). Plantation forests are often more simplified and intensively managed, but they may still contribute to biodiversity conservation by supporting some species and increasing habitat availability at the landscape scale (Barlow et al. 2007; Wang et al. 2022; Carnus et al. 2006). Therefore, understanding how biodiversity is maintained in secondary and plantation forests is essential for conservation planning, especially in regions where primary forests are scarce or fragmented.

Butterflies are suitable biodiversity indicators because they are sensitive to habitat disturbance, vegetation structure, and microclimate (Forsberg et al. 2020). Moreover, fruit-feeding butterflies can be sampled in a repeatable manner by non-experts using fruit-baited traps and then identified relatively easily. Therefore, they have been used widely in tropical regions, especially to study community structure in intact forest (Devries and Walla 2001), measure effects of disturbance (Barlow et al. 2007), and assess biodiversity of agroforestry (Francesconi et al. 2013) and even urban habitats (Oliveira et al. 2018).

In Vietnam, forests have experienced long-term pressure from deforestation, degradation, and land-use change (Duwe et al. 2022). To mitigate biodiversity loss and strengthen ecosystem services, forest regeneration and plantation development are increasingly important parts of conservation and land-management strategies (Cochard et al. 2016; Paudyal et al. 2020). Although biodiversity studies directly comparing degraded and plantation forests with primary forest remain limited in Vietnam, several studies have examined butterfly communities across gradients of habitat disturbance. For example, in Tam Dao National Park, butterfly diversity was highest in highly disturbed mixed habitats, whereas species of greatest conservation importance were associated with undisturbed or moderately disturbed forests (Lien and Yuan 2003). Subsequent work in Tam Dao similarly showed that habitat modification altered butterfly communities, with secondary forest supporting more forest-associated species than road-disturbed habitat (Vu et al. 2015). Studies in southern Vietnam also found marked differences in butterfly richness and community composition among natural, disturbed, bamboo, and stream-side forest habitats (Van Vu and Vu 2011). These studies depend on many hours of fieldwork by butterfly specialists, which makes them less comparable with other studies and hard to replicate at scale, hence our focus on trapping fruit-feeding butterflies. A study on fruit-feeding butterflies in Vietnam found the highest abundance in forest of intermediate disturbance, but some unique species were only found in the least disturbed parts (Bac and Bui 2023).

While comparing communities across habitats, temporal variation also needs to be taken into account. In seasonal tropical environments, butterfly abundance and community composition may change across the year in response to rainfall, temperature, humidity, host-plant phenology, adult food resources, and life-history cycles (Checa et al. 2009; Grøtan et al. 2014; Kishimoto-Yamada and Itioka 2015; Valtonen et al. 2013; Molleman 2018; Molleman et al. 2022). Surveys in northern Vietnam have also revealed substantial seasonal variation in butterfly richness, abundance, and community composition (Franzén et al. 2017). Temporal variation can therefore mask or exaggerate differences among sites if sampling is restricted to only one period, making repeated sampling necessary to distinguish habitat-related differences from seasonal fluctuations (Devries and Walla 2001; Hamer et al. 2005).

We performed a case study on the conservation value of a secondary and a plantation forest in northern Vietnam by comparing fruit-feeding butterfly community parameters with those in a primary forest. We sampled fruit-feeding butterflies nearly monthly during one year using banana-baited traps. We quantified how butterfly diversity and community composition differ between primary, secondary, and plantation forest. We also described how butterfly community composition and abundance changed over time. We finally discus to what extent the secondary and replantation forest can contribute to fruit-feeding butterfly conservation in northern Vietnam.

## METHODS

### Study area

This study was conducted in northern Vietnam at two sites: Cuc Phuong National Park and the Vietnam National University of Forestry campus in Xuan Mai, Hanoi. Cuc Phuong National Park was used to sample butterflies in primary forest and secondary forest, while the experimental forest at the Vietnam National University of Forestry represented the plantation forest. The two study sites were separated by a straight-line distance of approximately 100 km.

Cuc Phuong National Park is located in northern Vietnam, about 120 km southwest of the capital Hanoi, at the boundary of Ninh Binh, Hoa Binh, and Thanh Hoa provinces (20°14′–20°24′ N, 105°29′–105°44′ E). The park is a hilly limestone karst landscape, with forested mountains and valleys. Elevation ranges from about 150 m to 656 m a.s.l., and the vegetation is mainly tropical and subtropical forest growing on limestone (Duwe et al. 2022). The climate is humid tropical, with an average annual temperature of about 21°C and annual rainfall of around 2,100 mm. The dry season generally occurs from November to February, while the wetter period occurs during the rest of the year (Duwe et al. 2022; Thin 1997). The primary forest site and secondary forest transects were located 1 km from each other. The secondary forest has been left to regenerate after disturbance since 1992.

The plantation forest site was located at Nui Luot, within the Vietnam National University of Forestry campus in Xuan Mai Town, Chuong My District, Hanoi (20°51′31″ N, 105°30′45″ E). The Nui Luot area consists of low hills, with the highest elevations reaching about 133 m. The area has relatively homogeneous yellow-brown feralit soils developed on porphyrite parent material. The local climate is tropical monsoon, with two distinct seasons: a rainy season from mid-April to October and a dry season from November to March. Mean annual temperature is about 23.9°C, mean annual rainfall is approximately 1,677.7 mm, and mean annual relative humidity is 81.6%. The forest was planted in 1984 (40 years old) and covers approximately 27 ha. It forms an isolated forested hill surrounded by a predominantly human-modified landscape.

### Sampling of fruit-feeding butterflies

Butterflies were sampled nearly monthly for one year (October 2023 - October 2024). In the primary and secondary forests, we were unable to sample in January, July, and October 2024. In the plantation forest, we missed October 2023, June, August, September, and October 2024. At each forest type, a transect with 10 fixed trap stations was used with an inter-trap distance of at least 40 m. Butterflies were captured using banana-baited butterfly traps (125 cm high and 35 cm in diameter), hanging 40–50 cm above the ground (Molleman et al. 2006). Traps were baited with approximately 100 g of three-day-old fermented banana. Following bait placement, traps were monitored over three consecutive days during each sampling month, with checks conducted between 08:00 and 16:00. All butterflies were collected into envelopes labelled with transect, trap and date information. Butterflies were identified to the lowest possible level using morphological characters, including wing colour pattern, wing shape, and other diagnostic traits (Luong et al. 2004; Monastyrskii 2005, 2011; Monastryskii & Devyatkin 2015; Vu et al. 2026).

Climatic variables were included to describe seasonal environmental variation during the study. Daily rainfall and temperature data were obtained from the NASA POWER database using the nasapower package in R (Sparks 2018). Monthly rainfall totals were calculated by summing daily rainfall values.

#### Data analysis

All statistical analyses, data processing, and graphical visualizations were performed using R via the R Studio development environment (R Core Team 2025). Graphs were generated using ggplot2 (Wickham 2016). To gauge sample completeness and compare species diversity between the three sites, we constructed rarefaction curves. To visualize the overlap in species composition we made a Venn diagram showing the number of unique species for each site and the overlap between sites (Chen and Boutros 2011). To visualize the community composition by site and month while taking into account abundance, we performed a principal component analysis (PCA) using Hellinger-transformed species abundance data. We then tested for effects of site and month statistically using Redundancy Analysis (RDA) implemented in the vegan package (Oksanen et al. 2026). To account for temporal variation, month was modelled as a cyclical variable using sine and cosine terms calculated as sin(2π × month/12) and cos(2π × month/12). To determine whether seasonal patterns were driven by the same species in all habitats or by habitat-specific dominant taxa, we visually examined monthly abundance changes for the five most abundant species in each forest type.

## RESULTS

### Overview of the data

We caught 860 specimens belonging to 45 species from the three forest types. The rarefaction curve combining all forest types did not level off (Figure 1A). This suggests that with more sampling, significantly more species would be found, corroborating other studies in the region (Vu and Yuan 2003; Franzén et al. 2017; Monastyrskii 2011).

**FIGURE 1.**
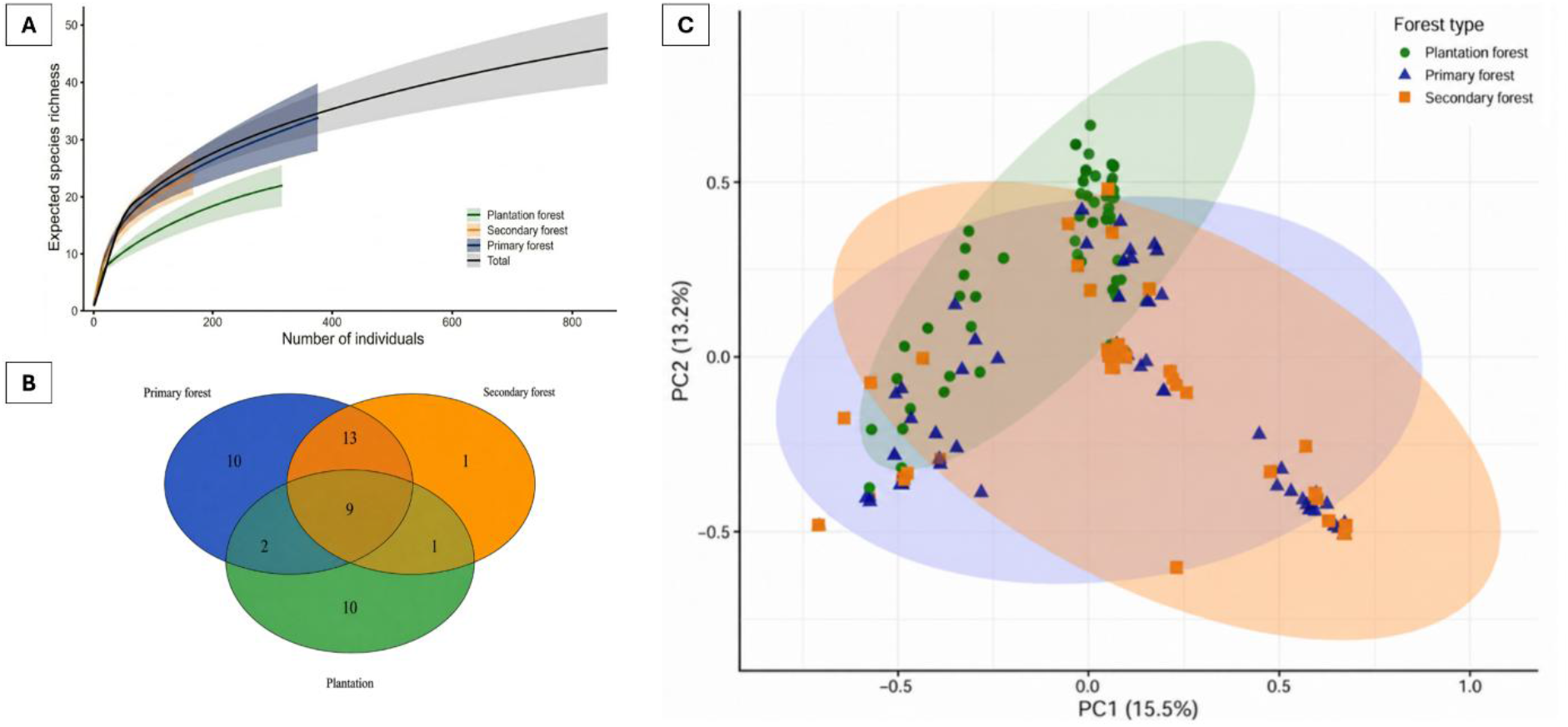
*For fruit-feeding butterflies recorded in Cuc Phuong National Park and Vietnam National University of Forestry* (A) S*pecies rarefaction curves per forest type and total, (B) a Venn diagram of the overlap in species composition among forest types, (C) PCA results of community composition. In the PCA plot, each point represents the butterfly community recorded from one trap in one sampling month, and shaded ellipses indicate the dispersion Hof samples within each forest type*.

### Fruit-feeding butterfly diversity differences between forest types

In the primary forest, we caught 376 specimens belonging to 33 species, while in the secondary forest we caught 167 specimens belonging to 24 species. Although the primary forest had more observed species, the overall species richness appears similar because the rarefaction curves for the primary and the secondary forest transects were very close and partly overlapping across the comparable range of sampling effort (Figure 1A). Thus, because fewer specimens were recorded in the secondary forest, its species richness may be underestimated more.

In the plantation forest, we caught 317 specimens belonging to 22 species. The plantation forest’s rarefaction curve was clearly lower and less steep than those of primary and secondary forests (Figure 1A), indicating that the plantation forest had a lower species richness.

### Fruit-feeding butterfly community composition

Nine species were shared by primary, secondary, and plantation forests: *Lethe confusa* (Aurivillius, 1898), *Melanitis leda* (Linnaeus, 1758), *Melanitis phedima* (Cramer, 1780), *Melanitis zitenius* (Herbst, 1796), *Mycalesis intermedia* (Moore, 1892), *Mycalesis malsara* (Moore,1858), *Mycalesis mineus* (Linnaeus, 1758), *Mycalesis mucianus* (Fruhstorfer, 1908), and *Tanaecia julii* (Lesson, 1837) (Figure 1B). Several of these were also among the most abundant species in our samples, suggesting that they are relatively widespread across the forest types studied rather than restricted to a particular habitat. Primary forest and plantation forest each contained 10 unique species, whereas secondary forest had only one unique species *Charaxes marmax* (Westwood, 1847). The strongest overlap was between primary and secondary forest, which shared 22 species, suggesting that their butterfly assemblages were more similar to each other than to plantation forest (Figure 1B).

Ten species were recorded exclusively in the plantation forest. These were *Charaxes bernardus* (Fabricius, 1793), *Cynitia whiteheadi* (Grose-Smith, 1899), *Elymnias patna* (Westwood, 1851), *Euripus nyctelius* (Doubleday, 1845), *Herona marathus* (Doubleday, 1848), *Hypolimnas bolina* (Linnaeus, 1758), *Lethe rohria* (Fabricius, 1787), *Mycalesis perseus* (Fabricius, 1775), *Mycalesis* sp., and *Symbrenthia lilaea* (Hewitson, 1864) (Table S1). Among these, *Herona marathus* (Doubleday, 1848) was also particularly abundant in the plantation forest.

Similar to species composition (Figure 1 B), the butterfly community in primary and secondary forest were largely overlapping, while the plantation forest was quite distinct (Figure 1C). Forest type explained a relatively small proportion of the total variation in community composition, with a R^2^ of 8% (Table 1), indicating that habitat-related differences were significant but moderate. Month also had a significant effect on community composition (Table 1), suggesting important temporal variation in butterfly assemblages. Specifically, the PCA analyses showed extensive variation in community composition within forest type (Figure 1C) that was in large part due to temporal variation (Table 1). Primary and secondary forests also showed greater dispersion than plantation forest (Figure 1C), indicating broader within-habitat variation in community composition.

**Table 1:** Results of RDA analysis on fruit-feeding butterflies recorded in Cuc Phuong National Park and Vietnam National University of Forestry. Samples were at the level of individual traps within each sampling month.

| Variable | Df | SoS | R<sup>2</sup> | F-value | P-value |
| --- | --- | --- | --- | --- | --- |
| Forest type | 2 | 0.0697 | 0.080 | 12.19 | < 0.001 |
| sinMonth | 1 | 0.0164 | 0.019 | 5.75 | < 0.001 |
| cosMonth | 1 | 0.0426 | 0.049 | 14.92 | < 0.001 |

### Seasonal dynamics of population abundance for five dominant species in each forest type

All nine common species exhibited clear temporal fluctuations in abundance. These species were *Mycalesis mucianus, Penthema darlisa* (Moore,1879), *Telinga malsara, Stichophthalma suffusa* (Leech,1892), *Stichophthalma fruhstorferi* (Röber,1903), *Melanitis leda, M. zitenius, M. phedima*, and *Herona marathus*. Within a species, the temporal fluctuations tended to be similar across sites, but with two exceptions: *T. malsara* and *M. mucianus* show an abundance peak in the primary forest without a concordant peak in the secondary or plantation forest (Figure 2). Across species, there appear to be some concordant peaks such as four species *Melanitis leda, M. zitenius, Herona marathus*, and *M. phedima* in the planation forest showing a peak abundance in December 2023, but synchrony does not appear strong overall.

**FIGURE 2.**
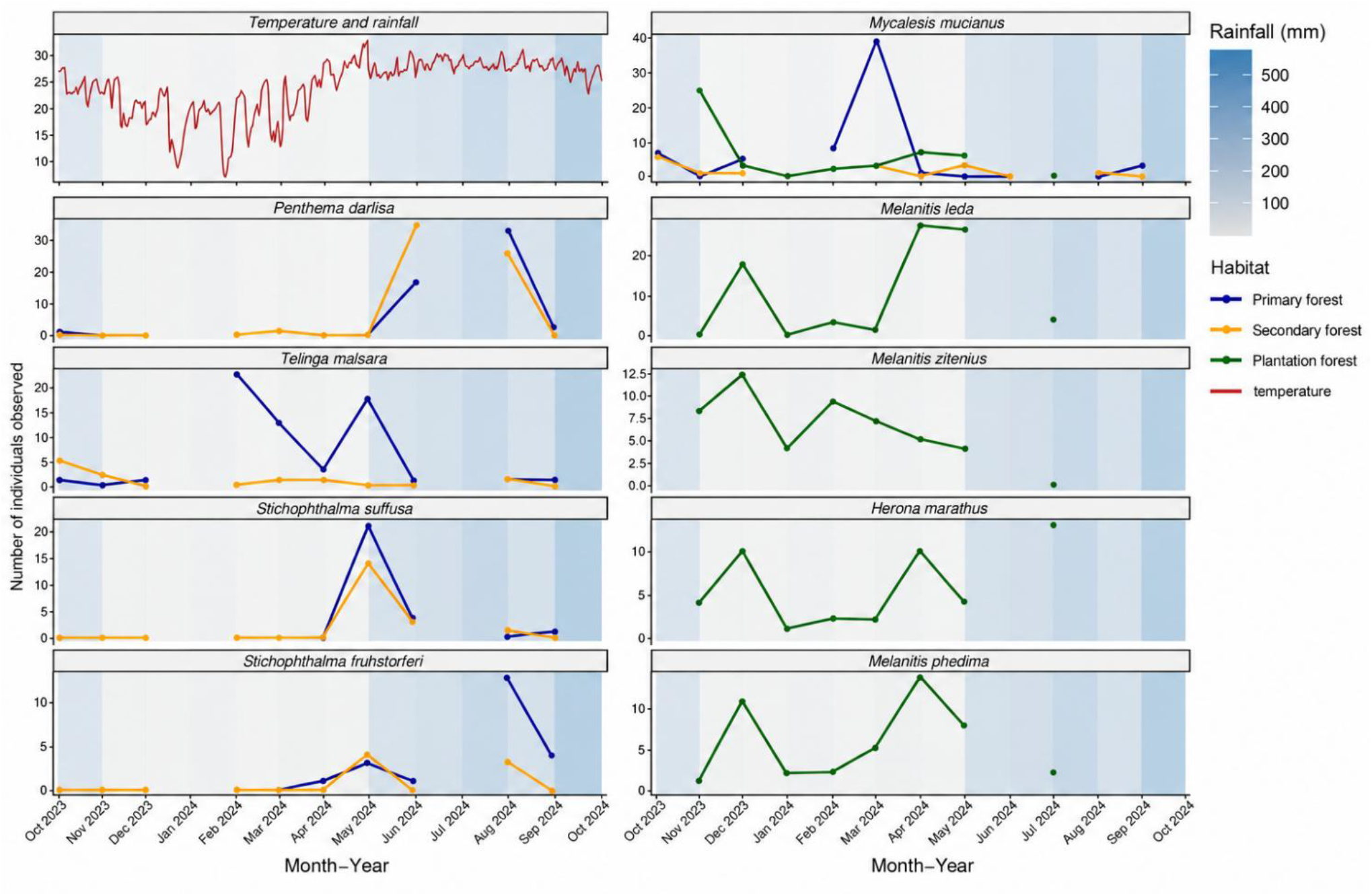
Seasonal population dynamics of dominant butterfly species across primary forest, secondary forest, and plantation habitats in northern Vietnam.

## DISCUSSION

The species richness was higher in primary forest than in the secondary one. This appears to contrast with previous studies that tend to find higher abundance and species richness in disturbed habitats (Barlow et al. 2007; Vu and Yuan 2003). However, this secondary forest has been regenerating since 1992, resulting in a dense canopy with limited understory vegetation, while the primary forest is more heterogeneous with canopy openings due to tree fall gaps. Such tree-fall gaps tend to be important habitat for understory butterflies and are also sites where canopy species can be encountered at understory level (Das et al. 2023; Hill et al. 2001; Pardonnet et al. 2013).

The secondary forest community appeared to be largely a subset of the species recorded in the primary forest, rather than representing a distinct disturbance-associated assemblage. This pattern is consistent with the capacity of regenerating forests to recover components of tropical butterfly communities, especially when they remain connected to less disturbed forest areas (Chazdon 2008; Korkiatupa et al. 2023). If secondary forest is protected and allowed to regenerate over a sufficient period, it may support the recovery of a diverse forest butterfly community, including many species associated with primary forest. Therefore, conserving secondary forest can help maintain biodiversity outside strictly primary forest areas and can improve habitat connectivity across the landscape.

In contrast, the plantation forest supported lower species richness and a more distinct butterfly assemblage, although it also contained several species not recorded in the other forest types. This indicates that plantation forest may favour species associated with more open, simplified, or disturbed habitats. Therefore, plantations can contribute to landscape-level diversity by hosting some disturbance-tolerant or habitat-specific species. However, their lower richness and distinct assemblage show that they have limited capacity to maintain the full diversity of fruit-feeding butterflies found in natural or regenerating forests, similar to plantation forest elsewhere (Barlow et al. 2007; Gibson et al. 2011). Nevertheless, similar to secondary forest, their community composition can be expected to become more similar to that of primary forest over time.

Butterfly abundance varied substantially through time, but the dominant species did not show a strong, synchronized seasonal pattern across all forest types. This suggests that abundance and activity of adult fruit-feeding butterflies is in part determined by common factors such as weather, fruit availability, and generalist predators and pathogens, but also by species-specific factors such as particular life-history traits or migration patterns, or specialized natural enemies such as parasitoids (Bonebrake et al. 2010; Molleman et al. 2022, 2016; Molleman 2018). Thus, the temporal patterns appears to be more similar to that in other wet tropical forests where dry seasons also do not tend to be severe (Grøtan et al. 2012, 2014; Valtonen et al. 2013), rather than to that of more seasonal climates where weather appears to be the main driving force (Das et al. 2023; Kishimoto-Yamada and Itioka 2015; Nobre et al. 2012).

Overall, the sampling was limited to one site per habitat and lacked canopy trapping. Therefore, this must be interpreted as a case study that may inform larger-scale studies that include replication, a range of secondary forest and planation forest types and ages, and canopy trapping.

Primary and secondary forests supported higher species richness than plantation forest, and their high species overlap indicates that secondary forest can retain an important component of the primary-forest butterfly fauna. This suggests that under effective protection and sufficient regeneration time, secondary forests may support the recovery of diverse forest butterfly assemblages. Although plantation forest contributed some unique species, it supported lower richness. Thus, secondary and plantation forests can likely contribute to butterfly conservation in human-modified landscapes, but they cannot replace primary forest. Conservation strategies should prioritize remaining primary forests while promoting secondary forest regeneration and biodiversity-oriented plantation management. Trapping of fruit-feeding butterflies would be an efficient part of monitoring the effectiveness of these conservation policies.

