## Supplement 1 for "Conservation value of secondary and plantation forests for fruit-feeding butterflies in northern Vietnam"

### Supporting Information

Table S1. The abundance of fruit-feeding butterflies per forest type with information about scientific and common names, author attribution, subfamily, tribe, subtribe, and totals. Species are arranged according to the taxonomic sequence used in Butterflies of Vietnam: An Illustrated Checklist (2015); Butterfly species (Lepidoptera: Rhopalocera) of Xuan Son National Park, Phu Tho province, Vietnam (Vu et al., 2026); current species names are retained where nomenclature differs. An em dash indicates that the checklist does not specify that rank.

| Species | Common name | Subfamily | Tribe | Subtribe | Primary forest | Secondary forest | Plantation forest | Total |
|---|---|---|---|---|---|---|---|---|
| *Athyma nefte* (Cramer, 1780) | Colour Sergeant | Limenitidinae | Limenitidini | Limenitidina | 1 | 0 | 0 | 1 |
| *Cethosia cyane* (Drury, 1773) | Leopard Lacewing | Heliconiinae | Acraeini | — | 4 | 2 | 0 | 6 |
| *Charaxes bernardus* (Fabricius, 1793) | Tawny Rajah | Charaxinae | Charaxini | — | 0 | 0 | 2 | 2 |
| *Charaxes marmax* (Westwood, 1847) | Yellow Rajah | Charaxinae | Charaxini | — | 0 | 1 | 0 | 1 |
| *Coelites nothis* (Westwood, 1850) | Scarce Catseye | Satyrinae | Satyrini | Eritina | 2 | 2 | 0 | 4 |
| *Cynitia whiteheadi* (Grose-Smith, 1899) | Green Skirt Baron | Limenitidinae | Limenitidini | Adoliadina | 0 | 0 | 2 | 2 |
| *Cyrestis cocles* (Fabricius, 1787) | Marbled Map | Biblidinae | Cyrestini | — | 1 | 0 | 0 | 1 |
| *Cyrestis themire* (Honrath, 1884) | Little Mapwing | Biblidinae | Cyrestini | — | 3 | 4 | 0 | 7 |
| *Discophora sondaica* (Boisduval, 1836) | Common Duffer | Amathusiinae | — | — | 0 | 2 | 2 | 4 |
| *Doleschallia bisaltide* (Cramer, 1777) | Autumn Leaf | Nymphalinae | Kallimini | — | 1 | 0 | 0 | 1 |
| *Elymnias malelas* (Hewitson, 1863) | Spotted Palmfly | Satyrinae | Elymniini | — | 1 | 1 | 0 | 2 |
| *Elymnias patna* (Westwood, 1851) | Blue-striped Palmfly | Satyrinae | Elymniini | — | 0 | 0 | 4 | 4 |
| *Euripus nyctelius* (Doubleday, 1845) | Courtesan | Apaturinae | — | — | 0 | 0 | 1 | 1 |
| *Euthalia monina* (Fabricius, 1787) | Powdered Baron | Limenitidinae | Limenitidini | Adoliadina | 1 | 2 | 0 | 3 |
| *Faunis canens* (Hübner, 1826) | Common Faun | Amathusiinae | — | — | 1 | 1 | 0 | 2 |
| *Herona marathus* (Doubleday, 1848) | Pasha | Apaturinae | — | — | 0 | 0 | 46 | 46 |
| *Hypolimnas bolina* (Linnaeus, 1758) | Great Eggfly | Nymphalinae | Kallimini | — | 0 | 0 | 1 | 1 |
| *Kallima incognita* Nakamura & Wakahara, 2013 | Broad-banded oakleaf | Nymphalinae | Kallimini | — | 1 | 0 | 0 | 1 |
| *Lethe confusa* (Aurivillius, 1898) | Banded Treebrown | Satyrinae | Satyrini | Lethina | 9 | 3 | 16 | 28 |
| *Lethe rohria* (Fabricius, 1787) | Common Treebrown | Satyrinae | Satyrini | Lethina | 0 | 0 | 1 | 1 |
| *Melanitis leda* (Linnaeus, 1758) | Common Evening Brown | Satyrinae | Melanitini | — | 16 | 8 | 81 | 105 |
| *Melanitis phedima* (Cramer, 1780) | Dark Evening Brown | Satyrinae | Melanitini | — | 16 | 1 | 45 | 62 |
| *Melanitis zitenius* (Herbst, 1796) | Great Evening Brown | Satyrinae | Melanitini | — | 19 | 5 | 49 | 73 |
| *Mycalesis deficiens* (Fruhstorfer, 1906) | — | Satyrinae | Satyrini | Mycalesina | 3 | 5 | 0 | 8 |
| *Mycalesis intermedia* (Moore, 1892) | Intermediate Bushbrown | Satyrinae | Satyrini | Mycalesina | 21 | 2 | 1 | 24 |
| *Mycalesis mineus* (Linnaeus, 1758) | Dark-branded Bushbrown | Satyrinae | Satyrini | Mycalesina | 8 | 1 | 2 | 11 |
| *Mycalesis mnasicles* (Hewitson, 1864) | Cyclops Bushbrown | Satyrinae | Satyrini | Mycalesina | 1 | 0 | 0 | 1 |
| *Mycalesis mucianus* (Fruhstorfer, 1908) | South China Bushbrown | Satyrinae | Satyrini | Mycalesina | 64 | 17 | 46 | 127 |
| *Mycalesis perseoides* (Moore, 1892) | Burmese Bushbrown | Satyrinae | Satyrini | Mycalesina | 8 | 4 | 0 | 12 |
| *Mycalesis perseus* (Fabricius, 1775) | Common Bushbrown | Satyrinae | Satyrini | Mycalesina | 0 | 0 | 1 | 1 |
| *Mycalesis* sp. | Bushbrown sp. | Satyrinae | Satyrini | Mycalesina | 0 | 0 | 2 | 2 |
| *Neope muirheadii* (C. & R. Felder, 1862) | Black-spotted Labyrinth | Satyrinae | Satyrini | Lethina | 9 | 0 | 2 | 11 |
| *Penthema darlisa* (Moore, 1879) | Blue Kaiser | Satyrinae | Satyrini | Lethina | 55 | 62 | 0 | 117 |
| *Ragadia cryptica cryptica* Nakamura & Lang, 2023 | Striped Ringlet | Satyrinae | Satyrini | Ragadiina | 1 | 0 | 0 | 1 |
| *Stibochiona nicea* (Gray, 1846) | Popinjay | Biblidinae | Pseudergolini | — | 1 | 0 | 1 | 2 |
| *Stichophthalma fruhstorferi* (Röber, 1903) | The Jungle Queen | Amathusiinae | — | — | 24 | 7 | 0 | 31 |
| *Stichophthalma fruhstorferi* Röber, 1903 | Jungle Queen | Amathusiinae | — | — | 2 | 1 | 0 | 3 |
| *Stichophthalma suffusa* (Leech, 1892) | Suffused Jungle Queen | Amathusiinae | — | — | 29 | 18 | 0 | 47 |
| *Symbrenthia lilaea* (Hewitson, 1864) | Common Jester | Nymphalinae | Nymphalini | — | 0 | 0 | 1 | 1 |
| *Tanaecia jahnu* (Moore, 1858) | Plain Earl | Limenitidinae | Limenitidini | Adoliadina | 1 | 0 | 0 | 1 |
| *Tanaecia julii* (Lesson, 1837) | Common Earl | Limenitidinae | Limenitidini | Adoliadina | 1 | 1 | 9 | 11 |
| *Mycalesis inopia* Fruhstorfer, 1908 | Watson's Bushbrown | Satyrinae | Satyrini | Mycalesina | 7 | 7 | 0 | 14 |
| *Mycalesis malsara* Moore, [1858] | White-line Bushbrown | Satyrinae | Satyrini | Mycalesina | 62 | 10 | 2 | 74 |
| *Terinos clarissa* (Boisduval, 1836) | Malayan Assyrian | Heliconiinae | Heliconiini | uncertain | 1 | 0 | 0 | 1 |
| *Thauria lathyi* (Fruhstorfer, 1902) | Jungle King | Amathusiinae | — | — | 2 | 0 | 0 | 2 |
| Total abundance | | | | | 376 | 167 | 317 | 860 |
| Number of taxa | | | | | 33 | 24 | 22 | 45 |
